# Impact of the expectation on memory reconsolidation using a post retrieval extinction paradigm

**DOI:** 10.1101/2020.10.26.354977

**Authors:** Julia Marinos, Andrea Ashbaugh

## Abstract

**Objective:** The present study examined if the expectation for learning enhances reconsolidation of conditioned fear memories using the post-retrieval extinction paradigm in an undergraduate sample (n = 48).

**Methods:** The study took place over three consecutive days. The expectation for learning was manipulated through oral instructions prior to memory reactivation. On day one, participants underwent differential fear conditioning to two spider images (CS+ and CS-). On day two, participants were assigned to either a reactivation with expectation for learning group, a reactivation with no expectation for learning group, or a no reactivation group. On day three, return of fear in response to the CS+ spider image was measured following reinstatement (i.e., four shocks). Fear potentiated startle (FPS) and skin conductance response (SCR) were taken as measures of fear.

**Results:** The study found evidence that the expectation for learning may enhanced reconsolidation with FPS as a measure of fear as it was only the expectation for learning group in which FPS to the CS+ remained stable following reinstatement, however this effect was small and non-robust. In contrast, no evidence of reconsolidation was observed for SCR, as all participants exhibited a return of fear following reinstatement.

**Implications:** These findings suggest that a verbal manipulation of the expectation for learning may not be salient enough to induce reconsolidation as measured by SCR but may be sufficient as measured by FPS. Additionally, given in the inconsistent findings between SCR and FPS, the study’s results bring into question whether the post-retrieval extinction paradigm is appropriate to investigate reconsolidation using both physiological measures concurrently.

## Introduction

Reconsolidation is the process where a long-term memory, once reactivated, returns to a malleable state and can be updated [1], strengthened [2], or blocked [3, 4]. Memory reconsolidation has been observed in a number of species [5, 6], including humans [4].

Studies in humans have consistently demonstrated that a conditioned fear response, as measured by fear potentiated startle (FPS), can be eliminated if reactivation is paired with oral administration of propranolol [4, 7, 8, 9], though a few studies have failed to replicate this effect [10, 11]. These findings have also been replicated for traumatic memories in individuals diagnosed with posttraumatic stress disorder [12, 13, 14]. Brunet et al., [12] demonstrated that physiological responding (i.e., heart rate and skin conductance responses (SCR)) to personalized script driven trauma imagery was reduced one week after the administration of propranolol post recall of the traumatic event compared to a placebo group. These findings were replicated in open-label studies [13, 14].

Additionally, reconsolidation has also been incorporated in treatment of the specific phobia of spiders using propranolol. Soeter and Kindt [9] examined if blocking reconsolidation in individuals with high levels of spider fear would inhibit the return of fear following reinstatement using a behavioural approach test (BAT) as a measure of fear. Participants who had their memory reactivated (i.e., by focusing on their fears) followed by an oral dose of propranolol displayed greater approach behaviour during a BAT than participants who received a placebo or did not have their memory reactivated. Differences between these groups were exhibited at all follow-up periods (i.e., 16 days, three months, and 1 year later) demonstrating that the reduction of fear following reconsolidation persisted long-term. These studies illustrate that reactivated memories following the administration of propranolol, can be modified and this process might be beneficial in the treatment of anxiety based psychological disorders.

However, there are a number of limitations to the use of propranolol to study memory reconsolidation. First, propranolol is a drug that modulates protein synthesis and as a result, may only indirectly target the mechanisms involved in reconsolidation [15, 16]. Second, research has found that individuals have a preference for behavioural interventions over drug therapy for the treatment of anxiety disorders [17]. This is important when considering the translational impact reconsolidation research can have for the treatment of anxiety disorders and posttraumatic stress disorder. Therefore, there is a need to examine memory reconsolidation using behavioural methods.

Schiller et al., [1] examined memory reconsolidation in a human sample using a behavioural design (i.e., Post-reterival extinction paradigam). The study found that participants who had their memory reactivated and then underwent extinction after a 10-minute delay period, did not display a conditioned fear response as measured by skin conductance following reinstatement. Conversely, participants who only underwent extinction or who had their memory reactivated and then underwent extinction outside the reconsolidation window (i.e., greater than 6 hours) exhibited a return of conditioned fear as measured by increased skin conductance following reinstatement 24 hours later. These results have been replicated [18, 19] however; several studies have failed to reproduce these results [4, 7, 8, 20]. The inconsistencies in replicating these findings suggest that recalling a memory is not always sufficant to reactivate a memory and therefore it is important to develop a better understanding of the conditions under which reconsolidation occurs.

Pharmacological blockade studies have examined whether reactivation needs to signal that something new can be learned (i.e., prediction error) for a memory to be rendered labile [21, 22]. Prediction error occurs when there is a violation between expectation and actual events and is believed to signal that something new can be learned [22]. Prediction error has been found to strengthen initial learning, as well as extinction learning [23] and may be important to facilitate memory reconsolidation [24].

Prediction error may play a role in reconsolidation as well. Using propanrolol, Sevenster et al., [22] examined if a prediction error was needed in order for a memory to return to a labile state and undergo reconsolidation. When there was a prediction error (i.e., reactivation signaled new information) the conditioned fear response did not return following reinstatement, as demonstrated by an elimination of the startle response. In contrast, when there was no prediction error (e.g., reactivation presented the exact same information as the previous day) fear did return following reinstatement as demonstrated by an elevated fear potentiated startle response following reinstatement. These findings suggest that when reactivation signals that something new can be learned (i.e., prediction error) the conditioned fear memory is more likely to be rendered labile and undergo reconsolidation.

Sevenster et al., [21] examined whether inducing the expectancy for learning can trigger reconsolidation by manipulating participants’ expectation to receive a shock prior to reactivation using propanrolol in humans. In the shock expectancy group, participants were connected to the shock equipment during reactivation but did not receive a shock as expected. Therefore, a mismatch occurred when the CS+ was presented in the absence of a shock. In the no shock expectancy group, there was no expectation to receive a shock when the CS+ was presented because participants were not connected to the shock equipment during reactivation. As expected, Sevenster et al., [21], found that the group that expected to receive a shock demonstrated reconsolidation, whereas the group that had no expectation for receiving the shock did not demonstrate reconsolidation. These studies illustrate that the expectation for learning during reactivation appears to be critical to the reconsolidation of conditioned fear memories using pharmacological methods.

Overall, research has demonstrated that recalling a memory is not sufficient for a memory to undergo reconsolidation [4, 7, 8, 20]. Pharmacological blockade studies have found that reactivation should indicate that something new can be learned in order for a memory to return to a malleable state and undergo reconsolidation [21, 22,]. However, studies utilizing behavioural experiments (i.e., post-retrieval extinction paradigm) have yet to investigate how the expectation for learning impacts reconsolidation. The expectation for learning during reactivation could be a boundary condition of memory reconsolidation and as a result, may explain why studies have struggled to consistently reconsolidate conditioned fear memories using purely behavioral methods [4, 7, 8, 20].

The purpose of the current study was to examine if the expectation for learning prior to memory reactivation impacted the reconsolidation process using the post-retrieval extinction paradigm. Participants were randomly assigned to a no-reactivation, a reactivation with expectation for learning, or a reactivation with no expectation for learning condition. The level of expectancy for learning was manipulated by providing each group with different instructions prior to reactivation. We predicted that participants that had their memory reactivated and expected to learn something new would not display a return of fear following reinstatement on day three (i.e., there would be no change in participants’ SCR or FPS response from the end of extinction on day two to the beginning of re-extinction following reinstatement on day three). Conversely, we predicted that the participants who did not expect to learn something new and participants that did not receive reactivation would demonstrate a return of fear on day 3 following reinstatement as demonstrated by increased SCR and FPS.

## Methods

All procedures were approved by the University of Ottawa’s Research Ethics Board and all participants provided informed consent.

### Participants

According to G*power [25], a total sample size of 42 was needed to achieve a power of .80 with an alpha of .05 and effect size of f = .25. Exclusion criteria included: a self-reported heart condition (e.g., heart transplant, artificial cardiac pacemakers; cardiac arrhythmias, uncontrolled hypo- or hypertension, myocardial infarction); or reported current use of a beta-blocker. We collected data until we had enough participants with usable physiological data. Ninety-nine undergraduate students from the University of Ottawa were recruited through an online participant pool run by the School of Psychology. Of the 99 participants, the following were excluded: 33 participants dropped out; six individuals were not invited back after day one because they could not identify which image was paired with the shock; and 12 were excluded because none of their physiological data was readable. The final sample consisted of 48 participants. Participants were compensated with one course credit at the end of day one and day two, for a total of two credits and received $5 for their participation on day three.

## Materials

### Conditioned stimuli

The conditioned stimuli (CS) consisted of two different images of spiders selected from the International Affective Picture System [IAPS images 1200 and 1201; 26]. The standardized conditioned stimuli were designed to study emotions and the selected IAPS images have been demonstrated to be emotionally arousing in a student population [26]. During fear conditioning on day one, one of the images was sometimes paired with the shock (CS+) and the other image was never paired with the shock (CS-). The image associated with CS+ and CS-was counterbalanced across participants.

### Unconditioned stimulus (US)

The US consisted of an electric shock. The shock was delivered by a Grass SD9 Square Pulse Stimulator via two disposable (3.81 × 2.54cm) Ag/AgCl sensors (pre-applied with 0% chloride wet gel) attached to the wrist of the dominant hand. Two 2-meter touchproof snap leads were attached to the sensors and the leads were plugged into the Grass SD9 Square Pulse Stimulator. Before testing, participants determined the level of shock used throughout the three days. The shock was administered starting at 10 volts and increased by 2.5 volts until the participant determined the shock was uncomfortable but not painful up to 60 volts. The same voltage was used on all three days of testing.

## Measures

### The Spider Phobia Questionnaire [SPQ; 27]

The SPQ is a 31-item self-report questionnaire measuring fear of spiders. Participants endorsed either true or false to indicate if each item applies to them. The SPQ has demonstrated acceptable test-retest reliability and discriminant validity in a student sample and can differentiate between participants with and without a specific phobia [28]. This measure was used to ensure level of spider fear was similar across all three groups. Cronbach’s alpha was α= .91, for the current sample.

### Spielberger State Trait Anxiety Inventory [29]

The STAI consists of two 20-item self-report questionnaires that assess trait (i.e., STAI-T) and state (i.e., STAI-S) anxiety. The STAI-T asks participants to rate how anxious they generally feel on a 4-point scale ranging from *almost never* (1) to *almost always* (4). The STAI-S asks participants to rate how anxious they feel right now on a 4-point scale ranging from not at all (1) to very much so (4). Both scales have demonstrated adequate convergent validity and excellent test-retest reliability [30]. This measure was used to assess if there were differences in trait and state anxiety levels between groups because high levels of anxiety have been demonstrated to impair reconsolidation [31]. Cronbach’s alpha for the STAI-S was α= .92 and the STAI-T was α= .92 for the current sample.

### Manipulation Check

Participates were asked on day 3 before debriefing to rate how much they were expecting to receive a shock at the beginning of day 2 on a scale of 0 (not at all) to 5 (very much) to determine if we were successful in manipulating the expectation for learning among the different conditions.

### Physiological measure

#### Skin conductance response

SCR was used as a measure of fear and emotional arousal [32]. SCR is primarily considered a measure of anxiety and is a direct measure of sympathetic activity, which is influenced by the stimulation of the behavioral inhibition system [33]. SCR is recommended as a measure of anxiety in situations where stimuli cannot be actively avoided.

SCR was measured throughout the study and was recorded with BioLab Acquisition Software 3.1.13 from MindWare Technologies Ltd. SCR was measured in micro-siemens and sampled constantly at 1000 Hz. The leads were connected to a 16-Channel Electrode Box and the signal was amplified with a Galvanic Skin Conductance Amplifier from Mindware Technologies Ltd. Participants had two disposable (3.81 × 2.54cm) Ag/AgCl sensors (pre-applied with 0% chloride wet gel) placed on the palm of their non-dominant hand on the thenar eminence and hypothenar eminence and two 2-meter touchproof snap leads were connected to the sensors. Leads were taped on the skin with hypoallergenic surgical tape to reduce movement which can interfere with the recording of the data. All physiological measures were recorded on a different computer than the visual stimuli to ensure the data is recorded with minimal impediment as the computer can overload when both applications are run and can create additional noise in the physiological data.

#### Fear potentiated startle

Fear potentiated startle was measured in accordance with the recommendations from the Committee report: Guidelines for human startle eyeblink electromyographic studies [34]. FPS measures the response to the conditioned stimuli by EMG surface measurement of the orbicularis oculi (i.e., muscle located under the lower eyelid that closes the eye during a blink). This measure is used to assess the startle response of the participant as neurologically it represents the connections from the amygdala to the startle-reflex pathway in the brainstem [35]. A loud white noise (40 msec; 104dB) was presented at each presentation of the CS through headphones (model AKG K92 closed back studio) to participants to elicit a startle response. EMG was measured throughout the study and was recorded with BioLab Acquisition Software 3.1.13 from MindWare Technologies Ltd. The leads were connected to a 16-Channel Electrode Box, after the skin surface was cleaned with alcohol. To assess the activity of the orbicularis oculi muscle, one electrode (diameter of 5mm Ag/Ag-Cl unshielded electrode filled with Signa Gel) was placed below the lower eyelid on the left side in line with the pupil when looking straight. A second electrode was placed about 1-2cm laterally from the first electrode. A third electrode acted as the isolated ground electrode and was placed on the forehead.

#### Procedure

Testing took place on three consecutive days each 24 hours apart. Participants were connected to the skin conductance, EMG, and shock electrodes at all time points throughout the study. On all days, testing sessions began with a 5-minute baseline of physiological measures followed by 10 habituation startle probes (i.e., loud white noise; 40 msec; 104dB).

#### Day One

Informed consent was obtained and participants completed all self-report measures. In all conditions, participants were connected to SCR, EMG, and shock electrodes. Following baseline and habituation, participants underwent fear conditioning. They received the following instructions prior to the start of fear conditioning: *“We are going to start. There will be two images presented. The shock will only be paired with one image. Monitor the relationship between the image you are seeing and when a shock is received. Please keep your eyes on the screen at all times.”* Participants underwent fear conditioning, which consisted of two presentations of the CS+, eight presentations of the CS-, and six presentations of the CS+ paired with the US in a pseudo random order. At the end of day one, participants were asked which image they learned was paired with the shock to ensure participants learned that the shock was only paired with one image. Participants were not invited back for the other two days of testing if they were not able to identify which image was paired with the shock (*n* = 6).

#### Day Two

Participants were randomly assigned to one of three groups: Expectation for Learning (*n*=16), No Expectation for Learning (*n*=16) or No Reactivation (*n*=16). In all conditions, participants were connected to SCR, EMG, and shock electrodes throughout testing. Baseline and FPS habituation trials were completed prior to randomization.

The Expectation for Learning and No Expectation for Learning Groups both underwent reactivation. Prior to reactivation, these two groups were given separate instructions on the screen and verbally by the experimenter to manipulate the expectation for learning. Participants in the Expectation for Learning condition were told: *“We are going to start. Shortly you will see the images you saw yesterday. The relationship between the shock and the images has changed. Please observe how it has changed. Please keep your eyes on the screen at all times.”*

Participants in the No Expectation for Learning condition were provided with the following instructions prior to reactivation: *“We are going to start. You will see the same images you saw yesterday. However, today you <u>WILL NEVER</u> receive a shock at any point during the experiment. Please keep your eyes on the screen at all times.”* Participants were provided with these instructions so they knew what to expect throughout testing and as a result minimize the expectation that something new could be learned. Participants in these two groups then had their memory reactivated via a single presentation of the CS+. Participants in these two groups then took a 10-minute break where they watched a TV show. This break allows for the activation of the neural mechanisms needed for reconsolidation to take place [1]. Participants in the No Reactivation condition were not exposed to the single presentation of the CS+ (i.e., their condition fear memory was not reactivated) and instead proceed straight to the 10-minute break.

Following the 10-minute break, all participants underwent extinction where the CS+ and the CS-were presented without the US. Before extinction, participants in all three conditions were provided with the following instructions: *“We are going to start. Please monitor the relationship between the image and when a shock is received. Please keep your eyes on the screen at all times. Are you ready?”* The number of presentations of the CS+ and CS-were equivalent across groups during extinction and for this reason, the no reactivation group received one additional CS+ during extinction. All groups received the same instructions prior to extinction.

#### Day Three

In all conditions, participants were connected to SCR, EMG, and shock electrodes. Following baseline and habituation EMG trials, participants underwent reinstatement, which consisted of four unpaired presentations of the US. Then participants took a 10-minute break and watched another clip of the TV show. Once the break was completed, participants underwent re-extinction, which consisted of presentations of the CS+ and the CS-without the US. Participants were then disconnected from the electrodes, debriefed, and compensated for their time.

Prior to debriefing, participants in the Expectation for Learning and No Expectation for Learning conditions were administered a manipulation check to determine if the level of expectancy to receive a shock prior to reactivation on day two was successfully manipulated. Participants rated how much they expected to receive a shock at the beginning of day two (0 = not at all to 5 = very much). Participants in the no reactivation condition were not administered the question because the level of expectancy to receive a shock was not manipulated on day two.

### Statistical analysis

Statistical analysis was performed using SPSS software version 23[36]. Statistical assumptions were violated and as a result, data was square root transformed to correct for violations of normality.

The early acquisition phase for SCR and FPS was calculated by taking the averages of trials two and three on day one. The first trial for both the CS+ and the CS-were disregarded to help reduce the impact of an orientating effect. The late acquisition phase for both SCR and FPS were calculated by taking the averages of trials seven and eight on day one.

For extinction, the early phase of extinction was calculated by taking the averages of trials one and two on day two. The late phase of extinction was calculated by taking the averages of trials 10 and 11 on day two. The late and early phase for extinction were calculated in the same way for SCR and FPS.

Initial analyses were conducted separately to examine if acquisition and extinction were successful. Separate ANOVAs and follow-up planned comparisons were computed for SCR and FPS for acquisition and extinction with Group (Expectation for Learning vs. No Expectation for Learning vs. No Reactivation) as the between-participant factor, and Stimulus (CS+ vs. CS-), and Time (Phase) as within participant factors.

To test our main analysis of interest, the effect of expectation of learning on reconsolidation, a mixed ANOVA was calculated with Group (No-Reactivation vs. Reactivation with Expectation for Learning vs. Reactivation with No Expectation For Learning) as the between-participant factor and Time (last trial of extinction on day 2 vs. first trial of re-extinction on day 3) and stimulus (CS+ vs. CS-) as the within-participant factors. The main analyses were calculated in the same way for SCR and FPS. It was expected that there would be a Stimulus by Time by Group interaction whereby participants in the reactivation with expectation for learning group would not show a return of fear, whereas the other two groups would exhibit a return of fear as demonstrated by an increase in SCR and FPS from the last trial of extinction to the first trial of re-extinction.

## Results

### Participant Characteristics

The final sample consisted of 48 participants with a mean age of 19.28 (*SD* = 1.86), 62% female. A one-way ANOVA revealed no differences between the groups with regards to age, *F* (2, 44) = 1.20, *p* = .31. A chi-squared revealed no differences with regards to gender, χ^2^ (3) = .67, *p* = .72. Table 1 provides a summary of participants’ scores on the anxiety and spider fear measures, as well as the mean voltage selected by participants. One-way ANOVAs found no differences between groups with regards to the SPQ, *F* (2, 44) = .14, *p* = .87, STAIT-T, *F* (2, 44) = .04, *p* = .96, or for the STAI-S, *F* (2, 44) = 2.29, *p* = .11.

**Table 1.** Participants means on the anxiety and spider fear measures

| Variable | Expectation for<br>learning group<br>M (SD) | No expectation for<br>learning group<br>M (SD) | No reactivation<br>M (SD) |
| --- | --- | --- | --- |
| Voltage | 36.50 (1.06) | 36.03 (10.97) | 40.50 (10.00) |
| SPQ | 9.05 (5.63) | 8.12 (6.36) | 7.80 (8.06) |
| STAI-T | 41.21 (9.83) | 41.82 (9.03) | 40.80 (11.03) |
| STAI-S | 40.53 (9.27) | 48.04 (10.96) | 42.53 (53) |

*Note.* M; Mean; SD; Standard deviation; SPQ; Spider Phobia Questionnaire (Klorman, Weerts, Hastings, Melamed, & Lang, 1974); STAI-T; Spielberger Trait Anxiety Inventory (Spielberger, Gorsuch, Lushene, Vagg, & Jacobs, 1983); STAI-S; Spielberger State Anxiety Inventory (Spielberger, Gorsuch, Lushene, Vagg, & Jacobs, 1983).

### Manipulation Check

An independent t-test was computed to determine if there were differences in ratings taken at the end of the study between the Expectation for Learning and No Expectation for Learning groups with regards to how much they expected to receive a shock on day two. We predicted that participants in the Expectation for Learning group would report a greater expectation to receive shock on day two. Participants in the Expectation for Learning condition did not differ (*M* = 4, *SD* = 1.48, min = 0 and max = 5), from the No Expectation for Learning condition (*M* = 3.81, *SD* = 1.38, min = 0 and max = 5), *t* (25) = .34, *p* = .74, d = .13, with regards to the degree to which they expected to receive a shock on day two.

### Skin Conductance

#### Acquisition

To establish that participants across the three groups underwent successful acquisition on day one, we conducted a Stimulus (CS+ vs. CS-) × Time (Start vs. End) × Group (Expectation for Learning vs. No Expectation for Learning vs. No Reactivation) mixed ANOVA. Figure 1 displays the mean of each trial during acquisition. We found main effects for Stimulus, *F* (1, 43) = 6.44, *p* = .02, *η*^*2*^_*p*_ = .13, and Time, *F* (1, 43) = 15.18, *p* = < .001, *η*^*2*^_*p*_ = .26, but no effect for Group, *F* (2, 43) = 1.27, *p* = .29, *η*^*2*^_*p*_ = .06. As expected, the main effects were moderated by a Stimulus × Time interaction, *F* (1, 43) = 6.58, *p* = .01, *η*^*2*^_*p*_ = .13. There were no other two- or three-way interactions, *Fs* (1, 43) < .74, *p*s > .48, *η*^*2*^_*p*_ *<* .03. As seen in Figure 1, there was no difference in the CS+ versus CS-at the start of acquisition, *t* (45) = -.31, *p* = .76, *d* = .05, but by the end of acquisition participants had higher SCRs in response to the CS+ than the CS-, *t* (45) = 3.13, *p* = .003, *d* = .48.

**Fig 1.**
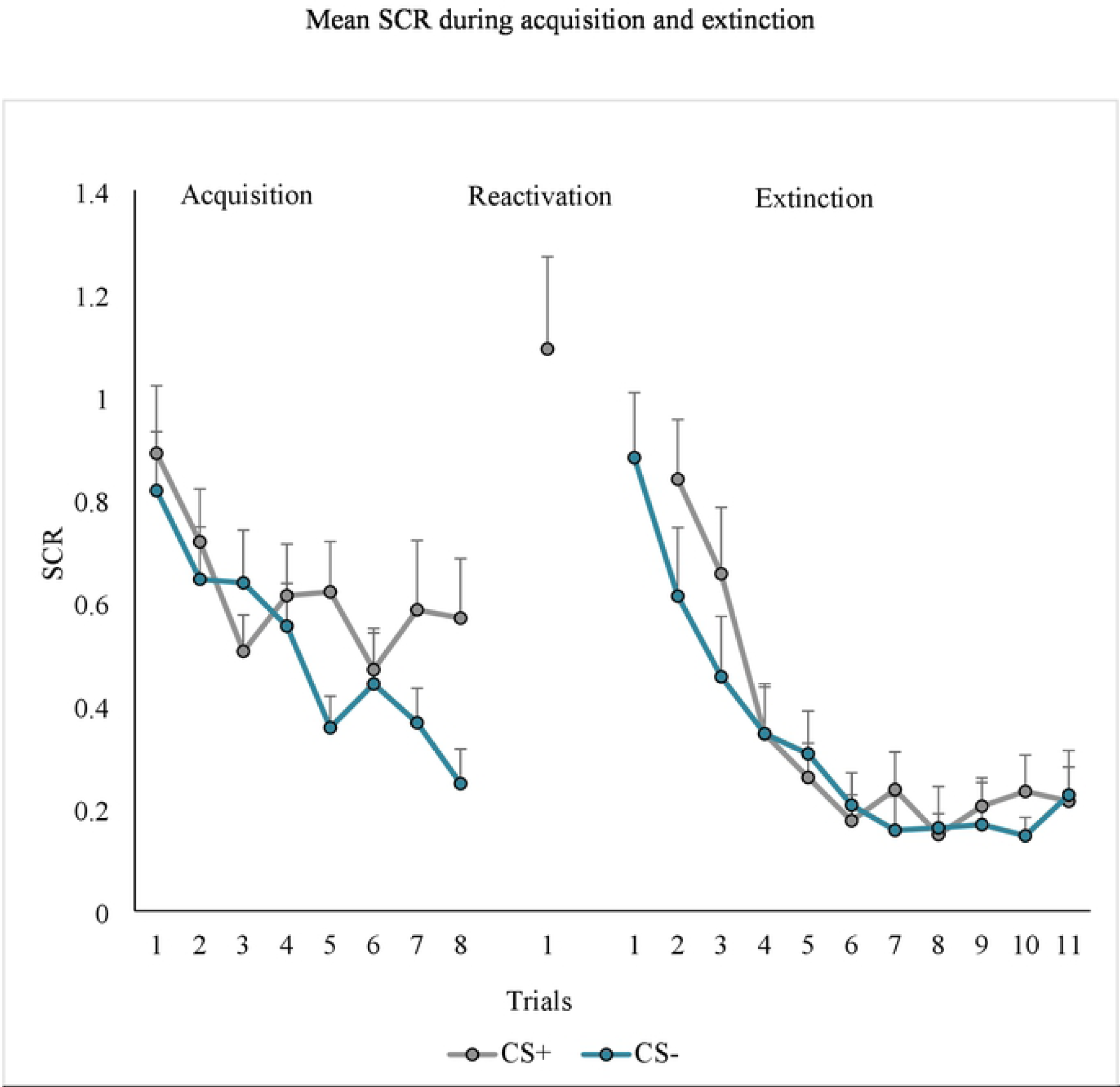
*Note*. Mean skin conductance response (SCR) across trials collapsed across groups. Acquisition consisted of presentations of the CSa+ that was sometimes paired with the US and the CS-that was never paired with the US. Extinction consisted of the presentation of the CSa+ and the CS-without the US. Measures of SCR were taken at every presentation of a stimuli.

#### Extinction

To assess if participants underwent successful extinction on day two, we conducted a Stimulus (CS+ vs. CS-) × Time (Day 2: Start vs. End) × Group (Expectation for Learning versus No Expectation for Learning vs. No Reactivation) mixed ANOVA. Figure 1 displays the mean of each trial during extinction. As expected, we found main effects of Stimulus, *F* (1, 43) = 6.32, *p* = .02, *η*^*2*^_*p*_ = .13, and Time, *F* (1, 43) = 52.06, *p* = <.000, *η*^*2*^_*p*_ = .55 but no main effect for Group *F* (2, 43) = .35, *p* = .71, *η*^*2*^_*p*_ = .02. None of the two-way or the three-way interactions were meaningful, *Fs* (1, 43) < 1.14, *p*s > .23, *η*^*2*^_*p*_ *<* .03. Given our a priori hypothesis (i.e., we predicted that participants would have a greater response to the CS+ than the CS-at the start of extinction and no difference in their response to the CS+ and CS-at the end), follow-up t-tests were calculated to compare the mean SCR response to the CS+ and the CS-at the start of extinction and at the end of extinction. As seen in Figure 1, participants had a greater SCR response to the CS+ than the CS-at the start of extinction, *t* (45) = 2.29, *p* = .03, *d* = .25, whereas there were no differences in response to the CS+ and CS-at the end of extinction, *t* (45) = .48, *p* = .63, *d* = .07.

#### Reconsolidation

To examine whether the expectation for learning prior to memory reactivation prevents reinstatement of the conditioned fear response, a Stimulus (CS+ vs. CS-) × Time (Last Trial of Extinction on Day 2 vs. First Trial of Re-extinction on Day 3) × Group (Expectation for Learning vs. No Expectation for Learning versus No Reactivation) mixed ANOVA was conducted. There was no main effects for Stimulus, *F* (1, 43) = .03, *p* = .87, *η*^*2*^_*p*_ *<* .01, or Group, *F* (2, 43) = .44, *p* = .65, *η*^*2*^_*p*_ = .02, but there was a main effect of Time, *F* (1, 43) = 42.18, *p* < .001, *η*^*2*^_*p*_ = .50. Contrary to predictions, there were no two-way or the three-way interactions, *Fs* (1, 43) < 1.39, *p*s >.26, *η*^*2*^_*p*_ *<* .04. However, given the main effect of time and our a priori hypothesis, separate follow-up t-tests were run to compare the return of fear in each of the groups (i.e., The last trial of extinction on day two compared to the first trial of re-extinction on day three). As seen in Figure 2, inconsistent with our predictions, participants in the Expectation for Learning condition showed an increase in their SCR to both the CS+, *t* (14) = -3.27, *p* = .01, *d* = -.89, and CS-, *t* (14) = -3.82, *p* = 002, *d* = -1.32, demonstrating a return of fear following reinstatement. This effect was also seen in the No Expectation for Learning condition, as participants showed an significant increase in their SCR to the CS+, *t* (14) = -2.56, *p* = .02, *d* = -.91, and the CS-, *t* (14) = -4.13 *p* = .001, *d* = -.90. The No Reactivation group showed an increase in their SCRs to the CS+, *t* (15) = -2.47, *p* = .03, *d* = -.84, but not to the CS-, *t* (15) = -1.86, *p* = .08, *d* = -.58, from the end of extinction on day two to the beginning of re-extinction on day three. Thus, participants, regardless of condition, showed a return of fear on day three. That is, we observed no evidence that reconsolidation took place, as measured by SCR.

**Fig 2.**
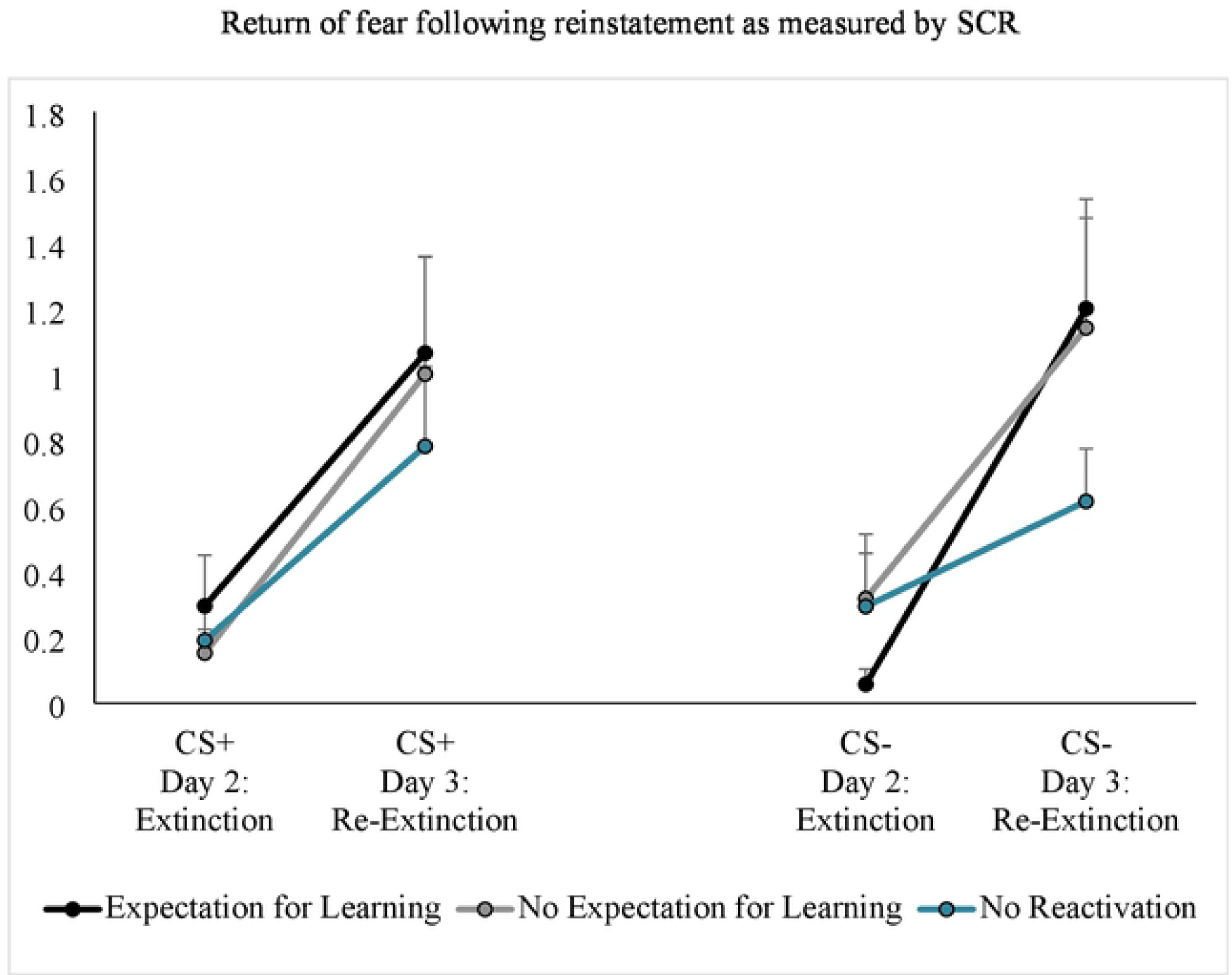
*Note*. Results for skin conductance (SCR) for reconsolidation. Day 2 is a measure of fear following extinction (i.e., last trial of extinction). Day 3 is a measure of fear following reinstatement of fear (i.e., first trial of re-extinction). Measures of SCR were taken at every presentation of a stimuli.

#### Fear Potentiated Startle

The same statistical analyses outlined above for SCR were computed for FPS.

#### Acquisition

Figure 3 depicts participants’ mean FPS response to each trial during acquisition. We found a main effect for Time, *F* (1, 45) = 25.77, *p* = < .001, *η*^*2*^_*p*_ = .36, and Stimulus, *F* (1, 45) = 5.56, *p* = .02, *η*^*2*^_*p*_ = .11, but no main effect for Group, *F* (2, 45) = 4.05, *p* = .02, *η*^*2*^_*p*_ = .15. None of the two-way interactions were meaningful, *Fs* (2, 45) < 2.59, *p*s > .09, *η*^*2*^_*p*_ *<* .10, however a Stimulus × Time × Group interaction was detected, *F* (2, 45) = 4.05, *p* = .02, *η*^*2*^_*p*_ = .15.

**Fig 3.**
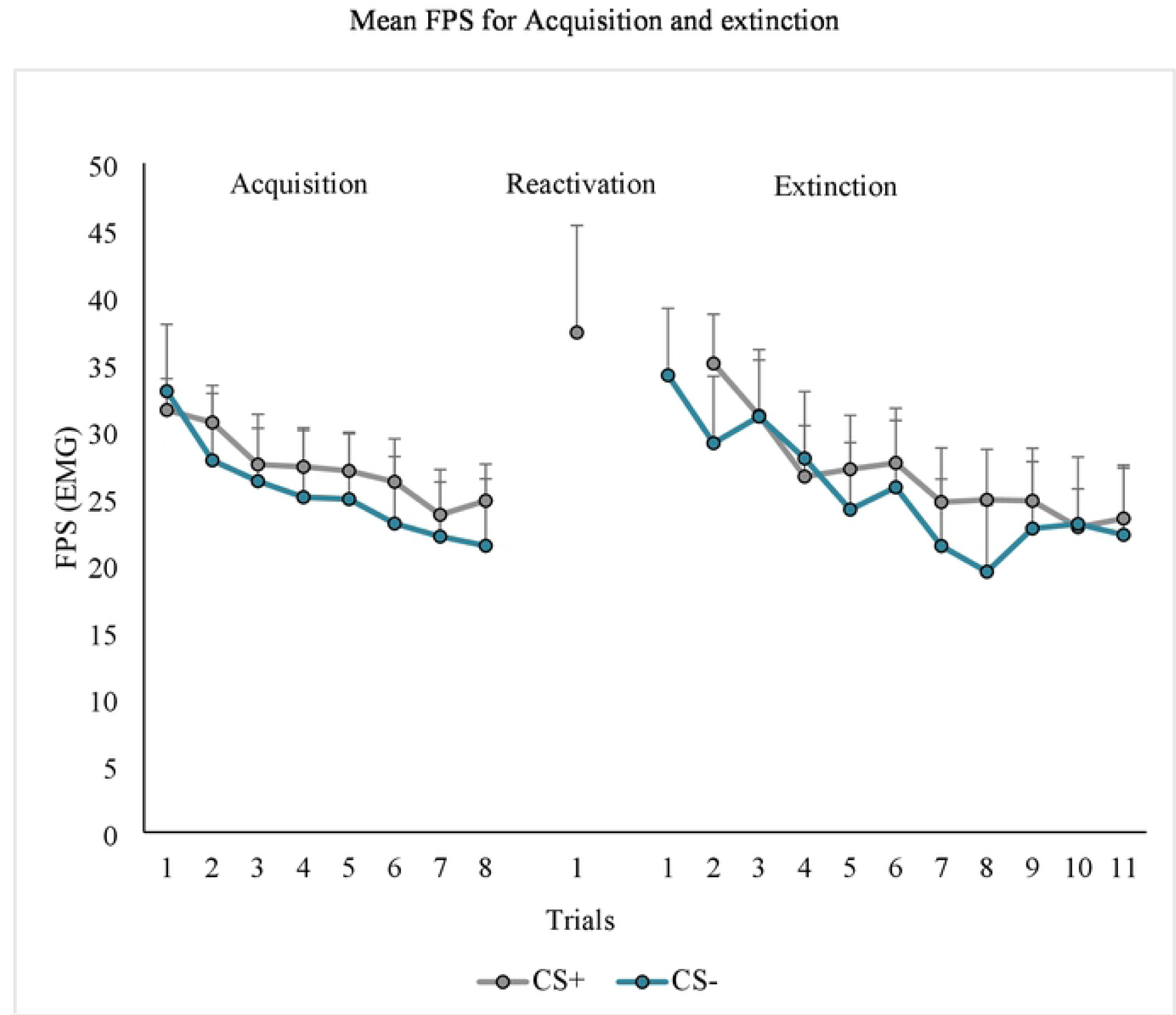
*Note*. Mean fear potentiated startle (FPS) across trials collapsed across groups. Acquisition consisted of presentations of the CSa+ that was sometimes paired with the US and the CS-that was never paired with the US. Extinction consisted of the presentation of the CSa+ and the CS-without the US. The error bars represent standard error. Measures of FPS were taken at every presentation of a stimuli.

Follow up analysis were run given the three-way interaction. For the Expectation for Learning Group, there were no differences in the CS+ compared to the CS-at the start, *t* (16) = .26, *p* = .80, *d* = .03, or the end of acquisition, *t* (15) = .72, *p* = .48, *d* = .11. For the No Expectation for Learning group, there was no difference in FPS to the CS+ compared to the CS-at the start of acquisition, *t* (15) = .49, *p* = .63, *d* = .06, but FPS was greater for CS+ than the CS-at the end of acquisition, *t* (15) = 2.37, *p* = .03, *d* = .4. For the No Reactivation group, participants showed greater FPS to the CS+ than the CS-, *t* (15) = 2.65, *p* = .02, *d* = .33, at the start of acquisition, but did not by the end, *t* (15) = -.19, *p* = .85, *d* = -.03. Thus, only the No Expectation for Learning condition displayed successful fear acquisition.

#### Extinction

Figure 3 shows participants’ mean FPS to each trial during extinction. As expected, we found main effects of Stimulus, *F* (1, 45) = 5.26, *p* = .03, *η*^*2*^ = .11, and Time, *F* (1, 45) = 33.61, *p* = < .001, *η*^*2*^_*p*_ = .43, but no main effect for Group, *F* (2, 45) = .001, *p* = .99, *η*^*2*^_*p*_ = < .001. None of the other two-way or the three-way interactions were meaningful, *Fs* (1, 45) < 1.84, *p*s > .17, *η*^*2*^_*p*_ *<* .08. We calculated follow-up *t*-tests comparing the mean FPS to the CS+ and the CS-at the start of extinction and at the end of extinction. Consistent with our expectations, participants had a larger FPS to the CS+ than the CS-at the start of extinction, *t* (47) = 2.13, *p* = .04, *d* = .16, but by the end of extinction there was no difference between the CS+ and CS-, *t* (45) = 1.09, *p* = .28, *d* = .07. These results suggest that extinction occurred in all groups, but the magnitude of this effect was not large.

#### Reconsolidation

Our 2 (Stimulus) × 2 (Time) × 3 (Group) ANOVA showed no main effects for Stimulus, *F* (1, 45) = 14.25, *p* = < .001, *η*^*2*^_*p*_ = .24, or Group, *F* (2, 45) = .22, *p* = .80, *η*^*2*^_*p*_ = .01, but there was a main effect of Time *F* (1, 45) = 14.25, *p* < .001, *η*^*2*^_*p*_ = .24. There was also a Stimulus × Group interaction, *F* (2, 45) = 3.57, *p* = .04, *η*^*2*^_*p*_ = .14. Inconsistent with our predictions, none of the other two-way or the three-way interactions were meaningful, *Fs* (1, 45) < 1.60, *p*s >.21, *η*^*2*^_*p*_ *<* .07. Given the main effect of Time, the two-way interaction of Stimulus by Group, and our a priori hypothesis, follow-up t-tests were calculated. As seen in Figure 4, participants in the Expectation for Learning group showed no difference in their FPS response to the CS+, *t* (15) = -1.20, *p* = .25, *d* = -.29 or CS-, *t* (15) = -.93, *p* = .37, *d* = -.25, from the last trial of extinction on day two to the first trial of re-extinction on day three, suggesting that there was no return of fear. Conversely, in the No Expectation for Learning condition participants had an increase in FPS response to the CS+, *t* (15) = -2.72, *p* = .02, *d* = -.75, and the CS-, *t* (15) = -3.27 *p* = .01, *d* = -.99, from the end of extinction of day two to the beginning of re-extinction following reinstatement on day three, suggesting that they did experience a return of fear. The No Reactivation group (i.e., extinction alone) also showed an increase in FPS from the end of extinction on day two to the beginning of re-extinction on day three to the CS+, *t* (15) = -3.76, *p* = .01, *d* = -.54, but not the CS-, *t* (15) = -1.59, *p* = .13, *d* = -.28, again suggesting that they also experienced a return of fear.

**Fig 4.**
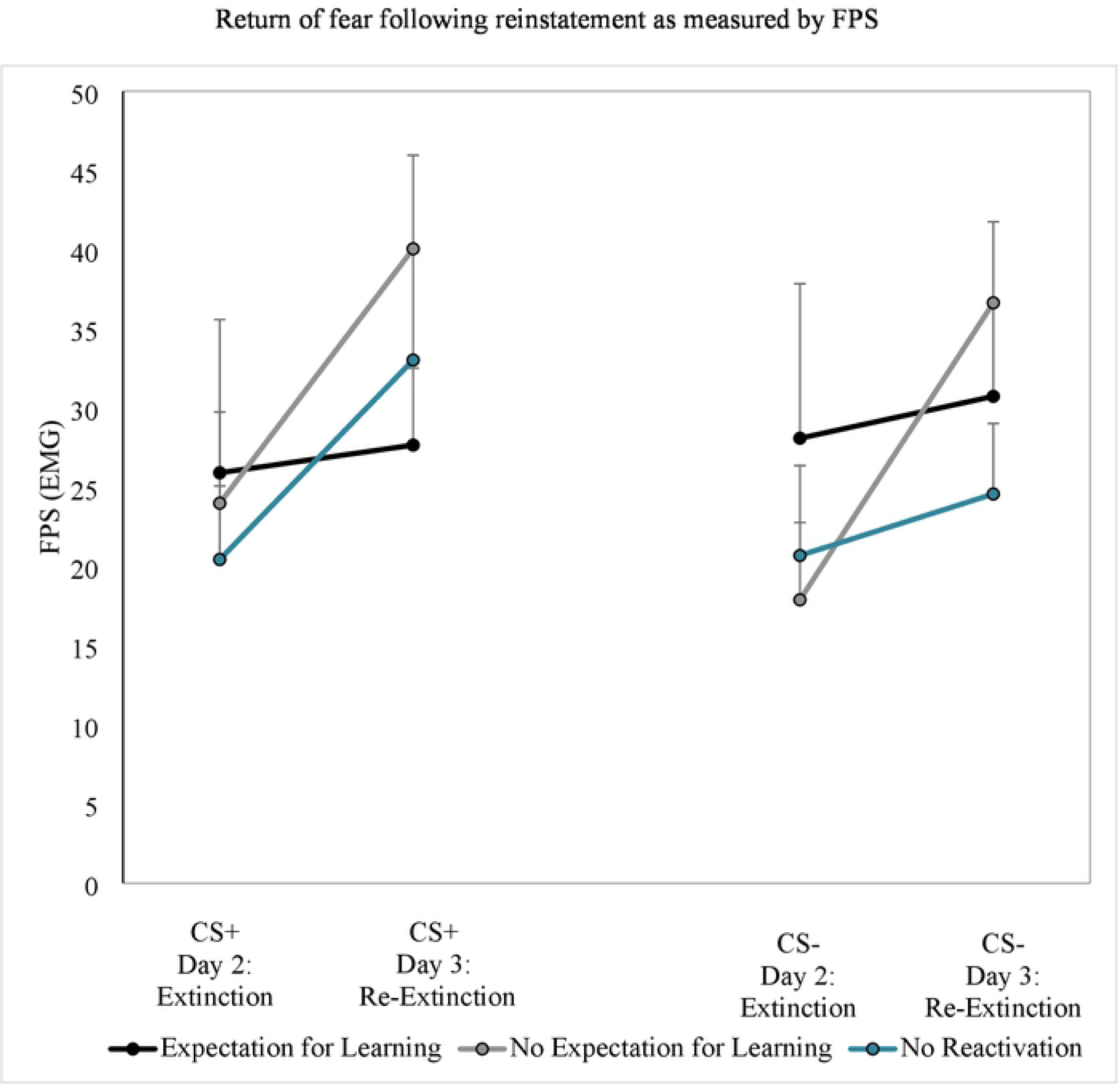
*Note*. Results for fear potentiated startle (FPS). Day 2 is a measure of fear following extinction (i.e., last trial of extinction). Day 3 is a measure of fear following reinstatement of fear (i.e., first trial of re-extinction). The error bars represent standard error. Measures of FPS were taken at every presentation of a stimuli.

#### Post hoc analysis

Given that not all groups displayed physiological fear acquisition with FPS as the measure of fear, we ran a post hoc analysis and included only participants that displayed greater FPS to CS+ than CS-during the late phase acquisition (n = 20). We calculated a Stimulus (CS+ vs. CS-) × Time (Last Trial of Extinction on Day 2 vs. First Trial of Re-extinction on Day 3) × Group (Expectation for Learning vs. No Expectation for Learning versus No Reactivation) mixed ANOVA to examine whether the expectation for learning prior to memory reactivation prevents reinstatement of the conditioned fear response. We found no meaningful effects for Stimulus, *F* (1, 17) = 2.00, *p* = .18, *η*^*2*^_*p*_ = .11, Time, *F* (1, 17) = 1.20, *p* = .29, *η*^*2*^_*p*_ = .07, or Group, *F* (2, 17) = .06, *p* = .94, *η*^*2*^_*p*_ = .01. None of the two- or three-way interactions had a meaningful *p* value, *Fs* (2, 17) < .2.71, *p*s > .10, *η*^*2*^_*p*_ *<* .24. The pattern of response was similar to that found in the entire sample. FPS response remained stable in the expectation for learning and the no expectation for learning groups from the end of extinction to the beginning of reinstatement. The no reactivation group displayed an increase in their FPS response following reinstatement.

## Discussion

The current study examined if the expectation for learning impacts the reconsolidation of conditioned fear memories using the post-retrieval extinction paradigm in an undergraduate sample using both SCR and FPS as measures of the fear response. The study found a disassociation between SCR and FPS. Specifically, for SCR, we found no evidence of reconsolidation as participants across all three conditions (i.e., the Expectation for Learning, the No Expectation for Learning, and the No Reactivation) exhibited a return of fear following reinstatement on day three of testing. In contrast, with FPS, we found evidence that increasing the expectation for learning enhanced reconsolidation, as it was only the Expectation for Learning group in which FPS to the CS+ remained stable following reinstatement. Participants in the No Expectation for Learning and No Reactivation showed an increase in FPS to the CS+, suggesting that fear returned in these conditions. However, it should be emphasized that this effect was small and non-robust. Furthermore, when we examined this only in participants that showed evidence of fear acquisition as measured by FPS, the effect disappeared.

The dissociation we found between the SCR and FPS is consistent with previous research [21, 4]. Researchers have suggested that SCR reflects a cognitive representation of arousal and is more difficult to reconsolidate [4, 7, 8]. In contrast, FPS is believed to represent an automatic emotional fear response and has been more successfully reconsolidated [7, 8, 4, 9]. It is notable that in studies where both SCR and FPS are measured, evidence of reconsolidation was found for FPS but not for SCR [4, 21], with the exception of one study which found that SCR did not interfere with the measurement of FPS [20]. Conversely, studies which only measure SCR have been more successful in demonstrating reconsolidation using SCR as a measure of fear [1, 19, 37].

FPS is frequently measured by using a loud startle probe and it is possible that this may interfere with the measurement of SCR. Previous research has found that shocks, unpleasant sounds, and a loud tone, similar to the one used in the current study to induce the startle response, are equally effective USs to produce fear acquisition in a classical conditioning paradigm using SCR as a measure of fear [38]. This is important because most studies that have examined reconsolidation using FPS as a measure of fear have utilized a loud tone to initiate startle response. It is possible that the loud tone used in the current study may be too aversive and as a result, may interfere with the measurement of the SCR because participants are anticipating the delivery of the loud tone. It is recommended that future research that measures both FPS and SCR consider using different methods to induce a FPS response.

The limitations of the study restrict the generalization and interpretation of the findings. First, participants did not exhibit successful physiological fear acquisition on day one with FPS as the measure of fear, as defined by greater FPS to the CS+ than the CS-. However, it should be noted that differences between the CS+ and CS-were in the expected direction, albeit small. Furthermore, by the start of day two, participants across all three groups did display a greater response to the CS+ than the CS-. This suggest that differential fear learning took place, but the effects were not apparent until day two.

Additionally, the results from the manipulation check found that the groups did not differ (i.e., the Expectation for Learning and the No Expectation for Learning groups) in their expectation to receive a shock on day two. Thus, it is possible that we did not successfully manipulate the expectancy to learn something new during reactivation. However, upon reflection, the question asked may not have accurately measured participants’ expectancy to learn something new but simply measured whether they expected to receive a shock during reactivation. Additionally, as the question was asked on day three (to ensure that the manipulation check did not interference with the actual experiment), it is possible that participants’ responses might not accurately represent their expectation to receive a shock on day two.

It is also possible that the verbal manipulation of expectancy for learning was not salient enough to differentiate the groups. In addition to verbal instructions, Sevenster et al., [21] maximized differences across groups by not connecting participants in the no expectation for learning group to the shock equipment to ensure that they absolutely could not expect to learn something new during reactivation. With the rationale of keeping groups similar in all respects, except for the expectation for learning, we chose to connect participants in the no expectation for learning group to the shock equipment and only provided verbal instructions that they would not receive a shock. It is possible that manipulating the expectation for learning needs to be more salient, and this could explain differences across these two studies. Future research should further explore how strong the expectation for learning needs to be to influence reconsolidation.

Despite these limitations, the findings from the current study have important implications. Our results suggest that recalling a memory is not sufficient for a memory to undergo reconsolidation. The present study demonstrated that a verbal manipulation of the expectation for learning is not sufficient to induce reconsolidation with SCR, but we found limited evidence that it may be for FPS as a measure of fear. Furthermore, the inconsistent results between SCR and FPS in the current study as well as in previous studies [4, 21] raise questions about measuring SCR and FPS concurrently. Future research should consider using alternative methods to induce FPS when measuring SCR concurrently or consider having separate conditions for each measure. Overall, the findings demonstrate how nuanced memory reconsolidation is and raises the critical question about whether the post-retrieval extinction paradigm is appropriate to study reconsolidation using physiological measures concurrently.

## Acknowledgements

This paper was funded by a grant from the Natural Sciences and Engineering Research Council (RGPIN-2015-05379) awarded to Dr. Ashbaugh. We thank Olivia Simioni for her help with data collection and extraction.

## Author Contributions

**Conceptualization**: Julia Marinos, Andrea Ashbaugh

**Data curation**: Julia Marinos

**Formal Analysis**: Julia Marinos

**Funding Acquisition**: Andrea Ashbaugh

**Methodology:** Julia Marinos, Andrea Ashbaugh

**Writing-original draft**: Julia Marinos

**Writing-review & editing**: Andrea Ashbaugh

